# Meta-Analysis of Allergy Genome-Wide Association Studies

**DOI:** 10.1101/2020.12.24.424368

**Authors:** Ronin Sharma

## Abstract

Allergies are complex conditions involving both environmental and genetic factors. The genetic basis underlying allergic disease is investigated through genetic association studies. Genome-wide association studies (GWAS) leverage sequenced data to identify genetic mutations, such as single-nucleotide polymorphisms (SNPs), associated with phenotypes of interest. Machine learning can be used to analyze large datasets and generate predictive models. In this study, several classification models were created to predict the significance level of SNPs associated with allergies. Summary statistics were obtained from the GWAS Catalog and combined from several studies. Biological features such as chromosomal location, base pair location, effect allele, and odds ratio were used to train the models. The models ranged from simple linear regressions to multi-layer neural networks. The final models reached accuracies of 80% and reflect the features that have the largest impact on a SNP’s association level.

## Background

In the United States, more than 50 million individuals suffer with allergies. Every year, there are about 200,000 emergency room visits due to accidental allergic responses. Approximately $25 billion is spent on these visits and treatments annually. Overall, allergic disease is the 6th leading cause of chronic illness in the U.S. and is on the rise^1^. An allergic reaction occurs when an individual’s immune system overreacts to a substance he or she comes into contact with. These substances are called allergens and they stimulate the immune response. Upon exposure to an allergen, the immune system produces antibodies, which bind to mast cells, causing the release of histamine. Histamine results in allergenic symptoms such as rashes, swelling, and coughing.

There are several proposed causes of allergies, but there is no single reason why someone is allergic to a specific substance. Some scientists suggest that individuals that are exposed to a wide range of allergens at a young age are less likely to develop allergies. Others say allergies are innate and that there is nothing we can do to prevent them^2^. Ideally, we will learn more about the causes of allergies, as the main concern is preventing potential life-threatening allergic reactions. Current research has suggested that genetics plays a role and that an individual’s genetic makeup can impact the predisposition of allergies^3^. Specifically, genetic mutations can increase the risk of having allergies. One common mutation that is analyzed is a single-nucleotide polymorphism (SNP). A SNP is a mutation at a single position (locus) in an individual’s genome. There are approximately 3 billion base pair locations in the human genome, which contain about 4-5 million SNPs. Scientists use complex sequencing technology to identify SNPs, and computational methods analyze these results.

One computational tool is a genome-wide association study (GWAS). GWAS aim to identify SNPs associated with a specific trait of interest, which is called a phenotype^4^. After these studies are completed, the analysis of the associations is a critical step. In some cases, individuals identify cohorts for association studies and then analyze those results independently. However, in the current era of big data, there are repositories that store the results of many association studies, which allows for large scale meta-analyses. These databases and repositories expedite the discovery of novel findings since researchers don’t need to perform entire GWAS, which includes sequencing many individuals’ genomes, but instead can simply aggregate the findings of several GWAS to improve statistical power.

One large repository is the GWAS Catalog^5^. It stores the results of many GWAS, grouped by phenotype. Overall, the Catalog hosts the results of 4788 publications, which include 216,255 SNP associations. Specifically related to allergic disease, there are 18 allergy phenotypes with a total of 1,556 SNP associations. These phenotypes include some of the most common allergy phenotypes such as peanut allergy, allergic rhinitis, atopic dermatitis, and atopy.

### Research Questions

- How does base pair location impact the degree of SNP association? For example, are SNPs located at the beginning of chromosomes (smaller base pair location) more likely to be associated with allergic disease?
- How do odds ratio, chromosomal location, and the effect allele impact a SNP’s association level? Can we use some of these characteristics to predict if a SNP will have a strong association?

Current research has suggested using some type of computational modeling to improve the identification of novel associations. I proposed that we should use the sequenced data and relevant biological information about each SNP to predict its degree of association. I believe that we can use several phenotypes to create a baseline classification model, and then this model can be used to aid in the discovery of new significant associations for other phenotypes, including newly discovered phenotypes.

This model has several benefits. It reduces the computational power of the extensive statistical analysis, since such analysis will no longer be required (other than when creating the initial model). A standardized baseline model is better than having scientists and researchers perform different statistical tests in their studies. Different tests can result in disagreements about whether an association is truly significant, so having the model set a standard threshold will prevent disputes about significance from occurring. Additionally, this model can easily be replicated for unrelated diseases, as the same model generation procedure can be used for a different initial dataset of the specified phenotypes. Furthermore, this model can be expanded to take an individual’s genome as an input, identify if the individual has a significant SNP association with some disease (based on the previously trained data, so no new GWAS is required for the individual), and then inform the individual to take the necessary precautions. This will be very helpful, especially in the field of allergic disease. For example, if the model predicts that the individual is allergic to eggs, then the individual will be informed not to eat eggs until they undergo extensive blood tests to verify if he or she has the allergy. This could prevent a potentially fatal allergic reaction. A counterargument states that everyone should be tested to see if they have an allergy. However, this is not feasible since there are many allergens, so we can’t test everyone for all the different allergens. Instead, we sequence the individual’s genome once, and then use different models (one for each phenotype) to predict whether he or she has that phenotype, and then perform tests for those predicted phenotypes. Ideally, the accuracy of the model can exceed 90%, but the model will still be very helpful at slightly lower accuracies, as there is no harm in predicting that the individual has the phenotype when they don’t have it. In this regard, the model can have false positives, but the percentage of false negatives needs to be minimized.

In my project, I created several different types of models. My focus was to use some features in the dataset to predict the degree of a SNP’s association, and I incorporated 18 allergy phenotypes into the analysis. Some models included using only one feature (base pair location), while others included four features (base pair location, odds ratio, chromosomal location, effect allele). The basic Linear and Logistic Regression Models had prediction accuracies lower than 60%, and even some of the more complicated models (Support Vector Machine, Neural Networks, etc.) were below 50%. However, the Random Forest Model had an average accuracy of 80%. This model can be applied to predict the severity of SNP associations for related allergy phenotypes. Additionally, I also learned about the impact of a SNP’s base pair location on its degree of association from these models. I found that it has a very limited impact, so SNPs located at the beginning and end of chromosomes don’t have significantly different association levels.

## Methodology and Results

I obtained the original datasets from the GWAS Catalog (Summary statistics were downloaded from the NHGRI-EBI GWAS Catalog in October 2020). The GWAS Catalog was founded by the National Human Genome Research Institute (NHGRI) in 2008. The catalog was created in response to the significant increase in genome-wide association studies (GWAS) published. It has been challenging to identify published GWAS to fit specific needs since there are so many studies and each study analyzes multiple phenotypes. There were limited resources that combined the results of all these studies. The GWAS Catalog is a large-scale database of associations that allows users to search for GWAS results and visualize them using several biological criteria. I started by identifying the traits related to allergic disease that the GWAS Catalog had data for. The traits included allergic rhinitis, atopic eczema, atopy, peanut allergy, along with several others. I performed several preprocessing steps (remove missing values, binarize values) to clean the dataset prior to generating models.

### Logistic Regression

First, I created a Logistic Regression Model to predict SNP association level based on discrete base pair location. My main goal was to create a model to predict this outcome using the features in the dataset. I started with the discrete base pair feature because of the correlation I observed while exploring the dataset (and because this was the first analysis that I preregistered).

I created discrete SNP association levels based on the p-value of each SNP. In an individual, there is a threshold used to separate genome-wide significant associations with non-genome-wide significant associations. SNPs with p-values less than the threshold are classified as genome-wide significant, while SNPs with p-values greater than the threshold are not. However, there is not a single threshold used for all studies. It varies depending on the total number of SNPs and the number of traits analyzed in the study. I slightly modified the common method for calculating the threshold since I analyzed the results of multiple studies: threshold. This resulted in a threshold of 1.5853E-12 for genome-wide significance.

I had to perform some preprocessing prior to creating the model. The initial dataset stored the p-values as continuous values, so I created a new column for the corresponding discrete value. It was 1 if the SNP’s p-value was genome-wide significant (less than the threshold) and 0 if it was not (greater than the threshold).

Figure 1 displays the distribution of SNPs for the two significance levels. The dataset had approximately 25% more not significant SNPs than genome-wide significant SNPs. This imbalance increases the likelihood of the model overfitting on the not significant SNPs, since the data is largely biased towards that outcome.

**Figure 1:**
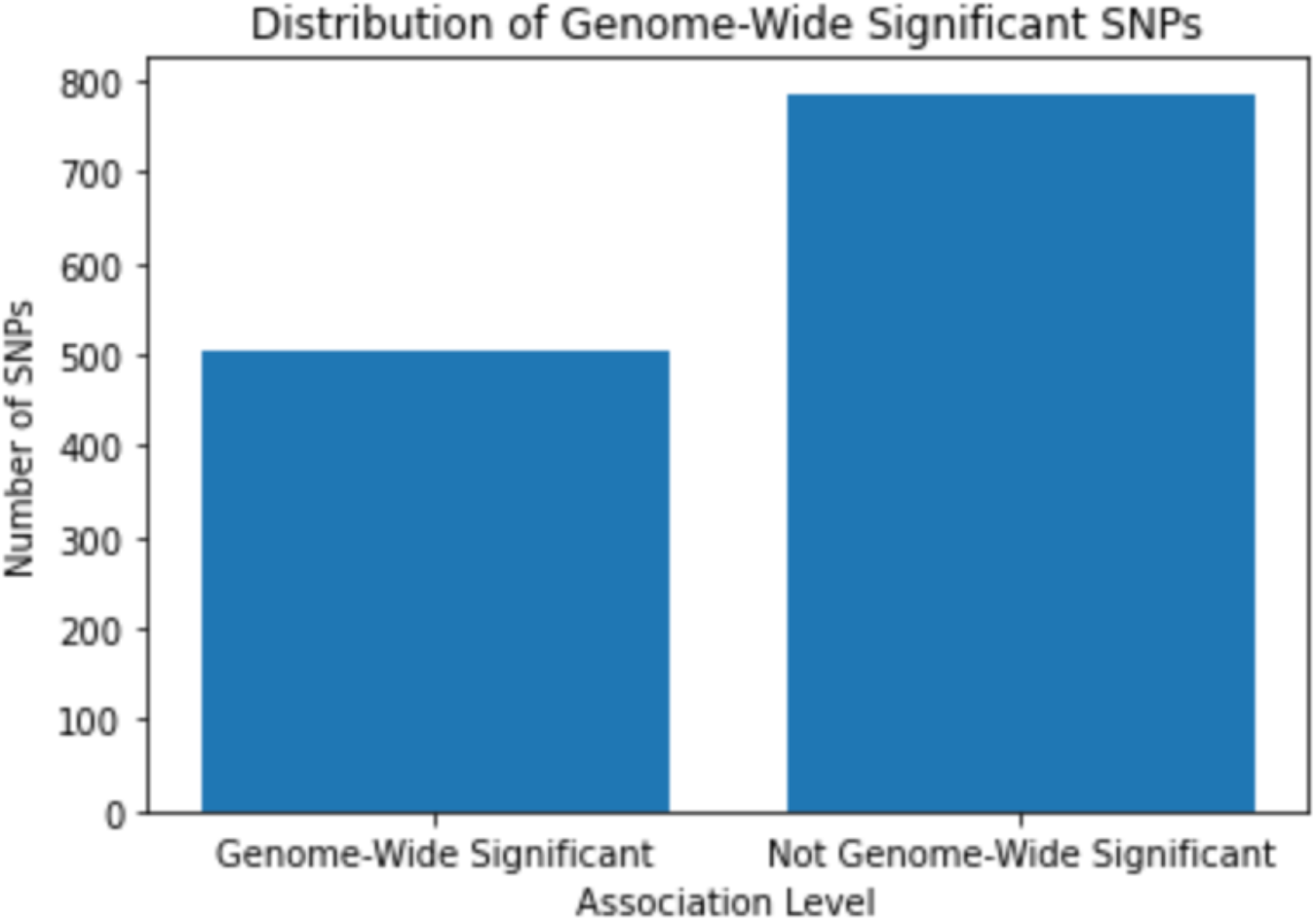
Distribution of Genome-Wide Significant SNPs

Next, I split the dataset into training and testing sets. I used a testing size of 30% in order to adequately evaluate the model’s performance. I fit the Logistic Regression model on the training data, and then evaluated the model. The model performed slightly better than average with an accuracy of 58%. The regression coefficient was negative, indicating that increasing the base pair (moving to a farther location on the chromosome) decreases the likelihood of the SNP reaching genome-wide significance, but the coefficient was very close to 0 so this relationship was very weak. Specifically, decreasing the discrete base pair location by 0.0708 would increase the discrete p-value by 1. There wasn’t a lot that I got from this coefficient, since the possible discrete base pair values were 1-4, so a change in 0.0708 wasn’t a change from one level to another. I had chosen not to use the continuous base pair values because the number of base pairs on each chromosome is different, so I wanted to normalize the location into four quantiles. However, this limited the conclusions I could draw from this model’s regression coefficient. Additionally, previous research had not identified a relationship between SNP significance level and chromosomal location^6^. In this regard, the small regression coefficient made sense since it supported previous findings.

I investigated potential reasons for the 58% accuracy, and one clear limitation of this model was that it always predicted 0. I plotted the model predictions over the range −100 to 100 (Figure 2). The model was only trained with inputs ranging from 1 to 4, so I wasn’t really interested in the prediction values outside this range. Ideally, the prediction transition from 0 to 1 would happen within the 1 to 4 range. Since it doesn’t, the model always predicted 0 or 1 (0 in this case), and thus only predicted about half correctly.

I tried modifying the train/test split sizes to see if this could possibly improve the model. I aimed to create a model that transitioned within the 1 to 4 range. Otherwise, the model would be fairly trivial as it only predicts one value for the range I’m interested in. I created models that ranged in test sizes from 0.1 to 0.5 to see if this impacted the transition range or accuracy.

**Figure 2:**
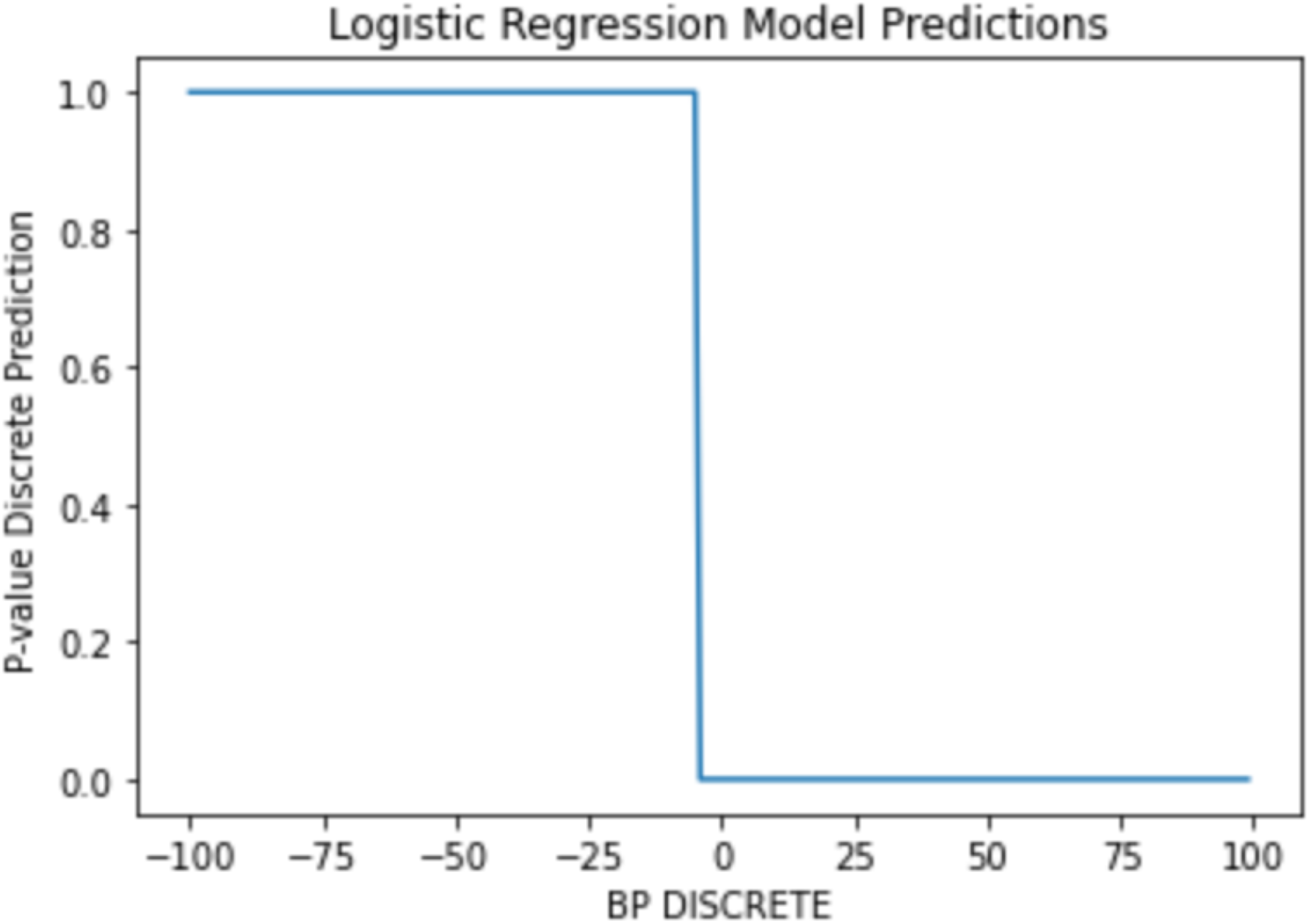
Logistic Regression Model Predictions

Figure 3 displays the accuracy score for all test/train splits. All models still only predicted 0 (never predict genome-wide significant for the input range I’m interested in), resulting in the maximum accuracy reaching only about 60%. Using a larger testing size did improve the accuracy, but this was marginal since it was only by 5%. Additionally, increasing the test size to 50% was not ideal because this significantly reduced the training size, resulting the the model not accurately learning the correct trend.

**Figure 3:**
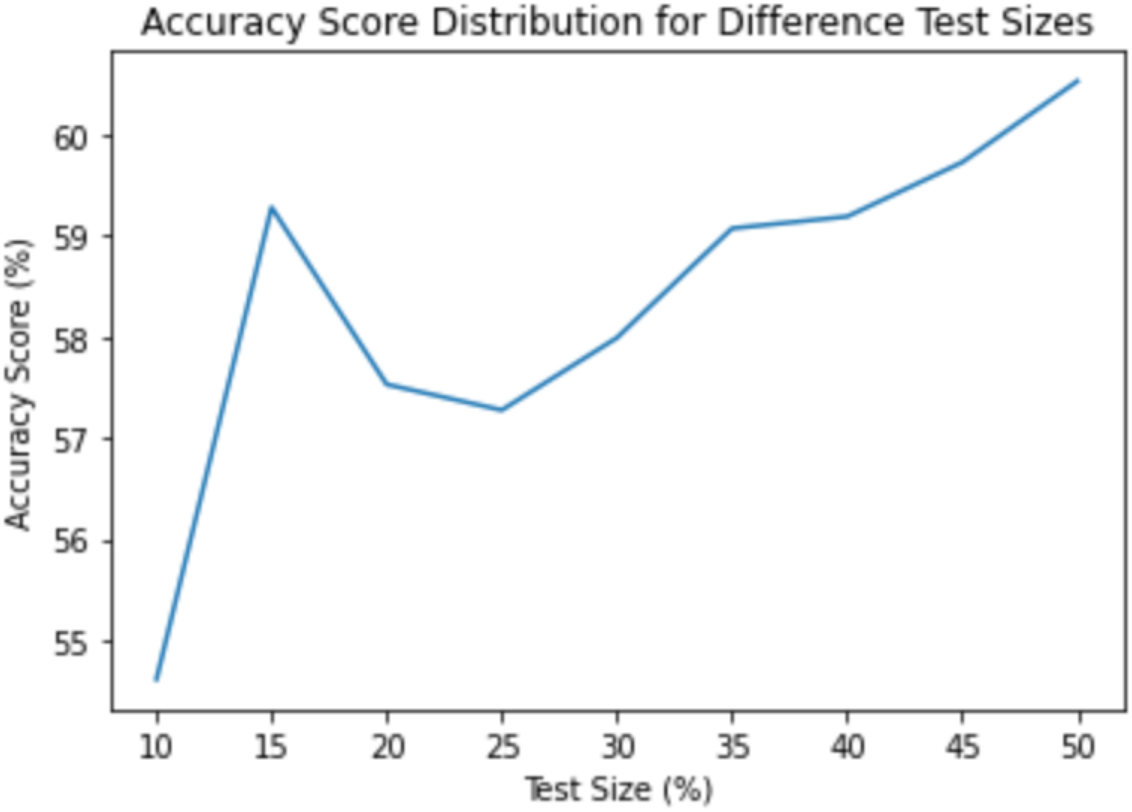
Accuracy Score Distribution for Different Test Sizes

This suggested that a Logistic Regression Model may not be the best model for this purpose. The main reason was that there didn’t seem to be a clear (and correct) distinction in inputs that led to a different output. For example, if the inputs 1 and 2 were more likely to be genome-wide significant (output 1), and the other inputs 3 and 4 were more likely to not to be significant (output 0), then a Logistic Regression Model would most likely work really well. However, that did not seem to be the case which suggested that we can’t broadly generalize location of a SNP on a chromosome into two classes and then determine if it’s significant based on the class it’s in. We most likely need to structure the classes differently as opposed to just having a threshold where values above are significant, and values below are not significant. For example, one possibility is that SNPs at the beginning and ends of chromosomes (inputs 1 and 4) are significant, while SNPs near the middle of chromosomes (inputs 2 and 3) are not significant. This would require more complicated models that can represent two transitions between 0 and 1, since Logit can only represent one. Additionally, this model was based on combining all the allergy phenotypes. Logistic regression may work better if I had fit the model to each individual phenotype. However, I had decided against this because of the overlapping phenotypes. In this dataset, the largest phenotypes combined multiple phenotypes (ex. the largest phenotype was ‘Eczema, allergic rhinitis’ (this limitation is further discussed in the Limitations section). There were only a few individual phenotypes (no overlapping), but these had very small sample sizes (<75 SNPs). Future analyses can combine additional datasets with more data for individual phenotypes to further analyze the use of Logistic Regression.

### Linear Regression

The next model I created was a Linear Regression Model to predict SNP association level based on odds ratio (OR), chromosomal location, discrete base pair location, and the discrete effect allele. The purpose for creating this model was the same as my purpose for creating the Logistic Regression Model. It aligned with my overall goal of predicting SNP association level using some, or all, of the dataset’s features. Logically, it made sense to start with a model that used fewer features (one feature for the Logistic Regression), and then move onto a model that used multiple features (this was also the second analysis that I preregistered).

Since I was not limited to a binary output, I expanded the number of possible SNP association levels from two to three. In addition to reaching genome-wide significance, SNPs can also reach suggestive association^7^. If SNPs don’t reach the suggestive association, then they are considered not significant. Suggestive association allows researchers to identify SNPs that have the potential to be associated with a phenotype, but they are not ready to conclude that it will have a definite association yet. The general workflow for interpreting SNP p-values is to first check if it reached genome-wide significance, and if not to check if it reached suggestive association. As a result, the suggestive association threshold is larger (less significant, larger p-value) than the genome-wide association threshold.

To calculate the threshold, I used the same formula. This resulted in the threshold of 1.5853E-9 for suggestive association. Additionally, I had to remove some SNPs with missing data. During the data exploration phase, I had decided not to remove all SNPs with missing data because that would have significantly reduced the size of the dataset. Instead, I chose to only remove the missing SNPs when I analyzed specific columns. So if columns X and Y had missing data, but in one analysis I was only using column X, I would only remove SNPs with missing values from column X. This allowed me to maximize the dataset for each analysis. For linear regression, the only column with missing values was the ‘EFFECT_ALLELE_DISCRETE’ column. Missing values were represented with −10, so I subsetted the dataset to remove those rows. This reduced the dataset from 1291 SNPs to 1098 SNPs.

Finally, I created a new column that stored the association level of each SNP. The value 2 corresponded to genome-wide significance, 1 corresponded to suggestive association, and 0 corresponded to not significant. Figure 4 displays the distribution of SNPs reaching each association level. Ideally, there would be an even split, with roughly the same number of SNPs in each category. I observed a distribution which was not exactly ideal, but it was close. The categories were about 100 SNPs apart (250, 350, and 450). The large number of genome-wide significant SNPs is a limitation of this research field, and research in general (the previous Logit distribution is uncommon since two discrete levels are rarely used). There is an emphasis placed on “significant” results, so they are reported more often. This is understandable since we learn a lot from “significant” results, but this introduces challenges when trying to understand the reasons for the significance. On the other hand, for this model I could have considered the genome-wide SNPs significant and the others not significant. If I did this, then there would have been more non-significant SNPs. However, I wanted to adjust for different degrees of association, so I included the third group (suggestive association).

**Figure 4:**
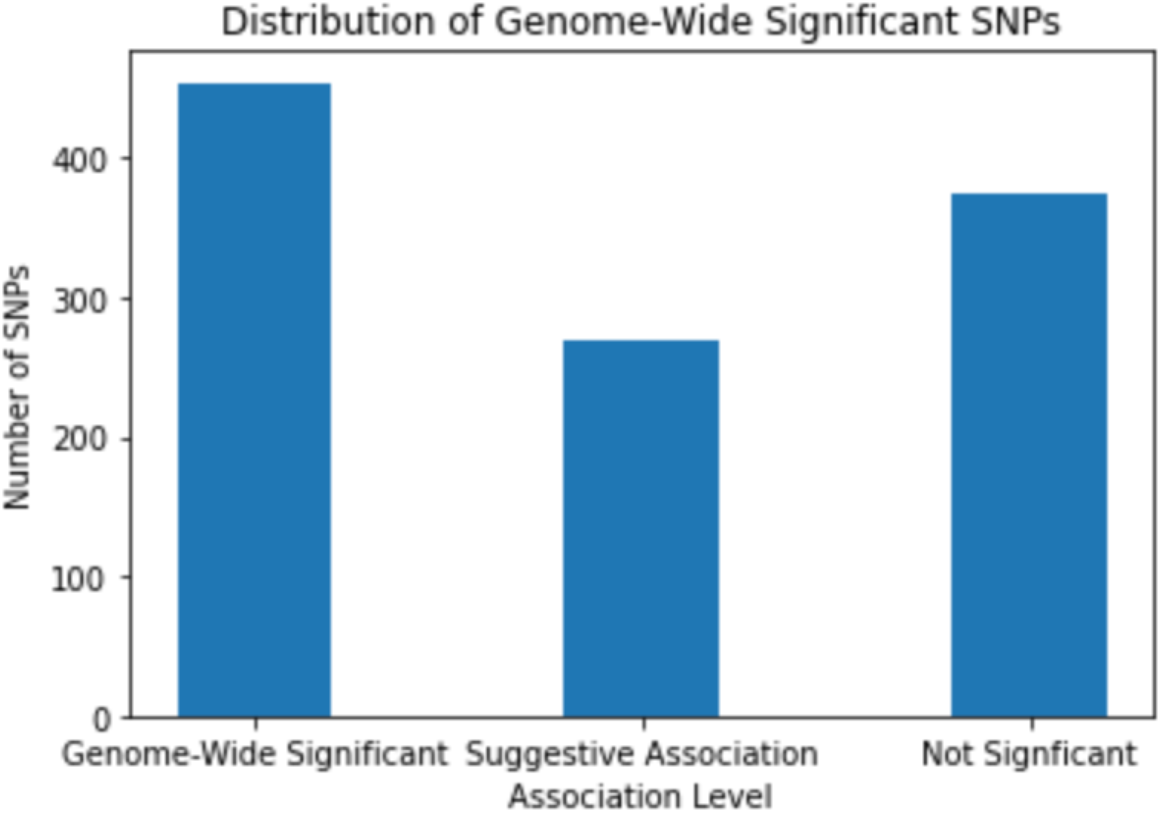
Distribution of Genome-Wide Significant SNPs

I created the model by first splitting the data into training and testing sets (test size = 30%). The model was terrible as it had an accuracy of 0%. The main reason was that Linear Regression Models are not well suited to predict discrete values. I chose not to keep the p-value continuous because those values were very specific and nuanced, which would lead to a lower accuracy (large negative orders of magnitude). Additionally, I am not interested in predicting the actual p-value, but whether the SNP reaches a specific threshold. This is because I don’t gain a lot of helpful information from knowing that a SNPs p-value is 1e-10 compared to 2e-10. However, it is helpful to know whether a SNP has reached genome-wide significance, as this indicates that we should definitely further analyze it.

Additionally, the main reason for the very low accuracy was that the model predictions got very close to 0, 1, or 2, but never exactly one of those values, thus they were considered incorrect. Out of all the regression coefficients for each feature, 3 out of 4 were very small, which suggests that those features don’t strongly impact the 3-level discrete p-value, and that the first feature, odds ratio, has the largest impact. However, this might have been the case because the features had different ranges and average values, so I tried to account for this in later analyses.

However, I still wanted to identify if a linear model could be used in some capacity. One alternative I experimented with was creating a hybrid model by using a Linear Regression model to predict a continuous p-value and then thresholding the result into one of the three association levels. To evaluate this hybrid model, I converted the predictions to 0, 1, or 2 based on the suggestive association and genome-wide significance thresholds. I also converted the “real” values for those predictions (y_test) to 0, 1, or 2 based on the same thresholds. The accuracy did improve to 37%, so a linear model is not useless for this dataset, but it definitely is not the best. This hybrid model performed about the same as a model that randomly chooses between three possible outcomes. This was not suitable as I didn’t want to make biological conclusions essentially by a random decision.

### Additional Models

With the Logistic Regression and Linear Regression Models, I achieved my goal of creating a model to predict SNP association level using the dataset’s features. However, the maximum accuracy was 58%, which is better than random chance but still not ideal. I shifted my focus to different models, with the goal of improving prediction accuracy.

A more complicated model might better predict the 3-level discrete p-value. The 2-level logistic model worked ok, but its accuracy was only 58%, and ideally, I wanted to be able to distinguish between genome-wide significant, suggestive association, and not significant, as opposed to just genome-wide significant and not genome-wide significant. This is because in many GWAS, there are not a lot of genome-wide significant SNPs. If I created a model that could only differentiate between SNPs that reached genome-wide significance and SNPs that didn’t, then I would classify many SNPs as “not significant” when they may have had some degree of association (suggestive association), and may have been worth further analyzing. In other words, a model that only predicts a 2-level discrete p-value can result in more false negatives, and a 3-level classification model can help reduce this.

I implemented the following models: Naive Bayes, Support Vector Machine (SVM), Decision Tree, Random Forest, k-Nearest Neighbors (kNN), and Neural Network (Multi-Layer Perceptron). Additionally, I performed boosting and stacking to try to improve the model, but this didn’t improve model accuracy. Table 1 displays a summary of the accuracy scores.

**Table 1:** Summary of Model Accuracies

| **Model** | **Accuracy Score** |
| --- | --- |
| Logistic Regression | 58% |
| Linear Regression | 0% |
| Linear Regression - Hybrid | 37% |
| Random Forest | 79% |
| Random Forest - Standardized | 42% |
| Random Forest - with PCA (95%) | 79% |
| k-Nearest Neighbors (n=1) | 78% |
| Gradient Boosting | 65% |
| AdaBoost | 55% |
| Stacked (Random Forest and kNN) | 56% |
| Support Vector Machine | 42% |
| Multi-Layer Perceptron Neural Network | 50% |

## Discussion

My goal was to create a model to predict the degree of a SNP’s association. I aimed to use a subset of the features in the dataset, and posed research questions about the potential impacts of specific features. The degree of a SNP’s association is indicated by its p-value, as p-values within certain ranges correspond to different degrees of association. The three degrees that I focused on were not significant, suggestive association, and genome-wide significance. I determined the exact values of these thresholds using the standard formula used in the field along with the relevant information (# SNPs/allergy studies) in the GWAS study. I created models that classified SNPs into either one of the three categories or just one of two (genome-wide significant/not significant). These models ranged from simple Linear Regression Models to complex Neural Networks. I explored the impact of data standardization and PCA on model accuracy. I observed a wide range of results, with one model having an accuracy of 0% and others having accuracies close to 80%.

One factor that I considered when evaluating the results was the impact of the data collection process and the preprocessing steps I performed. I don’t believe the way that I collected the data had any negative impacts. I used web scraping to obtain as much allergy data from the GWAS Catalog as possible, which was beneficial since a large dataset was ideal. The only limitation during this process was the actual data stored in the Catalog, since it didn’t have a lot of allergy data, especially compared to other diseases. During preprocessing, the only step that might have had an impact was when I removed some rows with missing entries. Intentionally, I only removed the rows which were almost missing all values in the row, because those rows were essentially unusable. For the rows that had some missing entries, I held off on removing them because I didn’t know which features (columns) I’d end up analyzing, and if the rows didn’t have missing values in the specific columns I was using, then there would be no need to remove them. Again, I don’t believe this had a negative impact, since it allowed me to maximize the size of the dataset for each analysis. Similar to the data collection process, this was more of a limitation of how the GWAS Catalog collected its data, and possibly the studies that reported the findings, which resulted in some missing values.

Table 1 displays the models’ accuracy scores. The score of each model indicates how well the model performed on the randomly generated test dataset. Each model was trained on some data (train set), and then its performance was evaluated using the test set. The model had never seen the data in the test set before. One important fact that I’d like to emphasize is that the percentage is the accuracy on the specific test set. This does not mean that the model will have the same accuracy when evaluated with other datasets. Ideally, the dataset that I trained the model with is representative such that the model performs the same (or better) on other datasets, but that is not always the case. Specifically, there were limitations with this dataset that caused this dataset not to be fully representative of allergic disease, and I discuss these in the next section. As a result, I can’t conclude that my Random Forest model, for instance, will always be 78.79% accurate. However, I can state that my Random Forest model was 78.79% accurate for the test set used and that I expect it to perform about the same on other datasets.

I did not expect to create a model that was 100% accurate. While I hoped to create models with high accuracies (above 90%), I was more interested in identifying if it is feasible to create models for this purpose and if the models can aid in the identification of significant SNPs. In this regard, establishing a viable model generation procedure was much more important than creating a model with a high accuracy. If I created a model that performed very well, then it would be great, but I could not apply it to lots of other datasets (due to limitations discussed in the next section). However, through experimenting with creating different models, I learned about models that work and don’t work on this allergy data. For instance, predicting continuous p-values, as opposed to one of two or three discrete levels, doesn’t work because the p-values are very specific (some on orders of 1e-8) so models will not be able to accurately predict that exact number often (indicated by 0% accuracy of the Linear Regression Model). Additionally, for this type of data, more complicated models do not always increase performance, which is also common in other fields. For example, the Multi-Layer Neural Network did not perform better than the much simpler Logistic Regression model. This suggests that the general allergy parameter space is not extremely complex such that it requires extensive use of complicated models like Neural Networks or other ensemble methods like boosting or stacking. There are simpler models such as Random Forest and kNN that can still achieve decent accuracies. Similarly, one procedure which is usually a very helpful preprocessing step before model generation is data standardization. However, that significantly reduced performance, as seen by the Random Forest Model. This suggests that even though all the columns don’t have the exact same ranges, it doesn’t impact the model. This makes sense because the features I chose don’t have significantly different ranges (1 to 4 vs. 1 to 23). On the other hand, I learned about models that worked well with this type of data. For example, Random Forest and kNN (n=1) both performed well. I believe that future studies should definitely implement these models with larger datasets. Additionally, PCA indicated that not all the features I initially considered are required for some models, and that we can use two of the four features and only witness a slight decrease in accuracy (<5%). This helps when analyzing much larger datasets, since we can essentially cut the training dataset sizes in half (column-wise), during the initial model generation process, and observe comparable results. Then, after fully evaluating the model, we can incorporate the entire training set to improve the performance slightly. Of course, after including the full dataset we would still evaluate the performance to ensure the model performed better, and reassess the model’s features and assumptions if it performed worse.

One of my research questions was about understanding the impact of base pair location (on a chromosome) on the degree of SNP association. The small regression coefficient of −0.0708 in the Logistic Regression model indicates that there is a very weak to no correlation between them. This supports previous findings that base pair location doesn’t have any major impact on a SNP’s degree of association, and suggests that this feature isn’t very important (or at least for this type of model). My other question was about the impact of some other features (OR, chromosomal location, effect allele) on a SNP’s association level, and whether we could use these features in a predictive model. Based on my results, I definitely believe that we can since some of the models performed fairly well. If we could not use these features, then I believe the models would have performed much worse, but since they didn’t, these features do have a non-negligible impact on a SNP’s association level, and should definitely be incorporated in future models.

Next steps include identifying larger datasets so that I can develop a more representative model. Specifically, I would like to obtain larger datasets for individual allergy phenotypes. Different phenotypes can have different feature trends, so there should be a model for each phenotype. I attempted to create phenotype-specific models, but the results were largely skewed by the small sample sizes and overlapping traits. Now, I can easily use the procedures that worked in this project, and apply them on more representative datasets. Overall, the goal is to further develop these types of models to use an individual’s sequenced genome as an input, and output whether that individual is likely to have an allergy to a certain phenotype. This will allow doctors to recommend individuals not to expose themselves to certain allergens until thorough testing is performed to verify if he or she has the allergy. This will help prevent potentially fatal allergic reactions and help the millions of individuals who suffer with allergies everyday.

## Acknowledgements

I would like to thank the GWAS Catalog for being open source, which allowed me to extract lots of data from the repository.

